# A novel exosome biogenesis mechanism: multivesicular structures budding and rupturing at the plasma membrane

**DOI:** 10.1101/849828

**Authors:** Stephanie Leon Quinonez, Ian R. Brown, Helen E. Grimsley, Jindrich Cinatl, Martin Michaelis, Chieh Hsu

## Abstract

Exosomes are small vesicles secreted by the cells, which mediate intercellular signalling and systemic physiological processes. Exosomes are known to originate from the intraluminal vesicles of the multivesicular endosome that fuses with the plasma membrane. We found that the non-small cell lung cancer (NSCLC) cell lines, HCC15 and A549, secreted exosomes with typical morphology and protein contents. Unexpectedly, transmission electron microscopy images indicated that the cells formed multivesicular structures that protruded from the plasma membrane and ruptured to release the exosomes. There were smooth multivesicular structures surrounded by an ordinary looking membrane, multivesicular structures coated by an electron dense layer with regular spacing pattern, and intermediate forms that combined elements of both. Electron microscopy images suggested that exosomes are release from these structures by burst events and not by the conventional fusion process. The molecular details of this novel mechanism for membrane association, deformation and fusion is to be unveiled in the future.

## Manuscript

Exosomes are small vesicles (30-100 nm) secreted by cells, which mediate intercellular signalling. They are involved in the regulation of physiological processes, e.g. during development and immune response. Aberrant exosome formation and signalling may contribute to disease, including neurodegenerative disease and cancer^1^.

Exosomes are known to originate from the endomembrane system. The multivesicular endosome, which contains intraluminal vesicles, fuses with the plasma membrane and release the intraluminal vesicles as exosomes. Merging of the multivesicular endosome and the plasma membrane involves Rab and SNARE proteins, similar to the membrane fusion events in exocytosis^2^. Here, we observed an unexpected novel exosome release process in non-small cell lung cancer (NSCLC) cells. Instead of fusing with the plasma membrane, multivesicular structures form bulges, which rupture to release the exosomes.

We collected exosomes from NSCLC HCC15 cells by differential centrifugation from foetal bovine serum (FBS)-free medium. The exosomes were within the typical exosome size range (**Fig. 1A,** see Supplementary Information for detailed methods). The known exosomal markers, CD63, flotillin-1, and Hsp70, were enriched in the HCC15 exosomes, while tubulin, GAPDH and GM130, cellular proteins not expected to be present in exosomes, were not detected. A lack of Arf6 (**Fig. 1B**), a marker of larger extracellular vesicles (microvesicles)^2^, further confirmed the identity of the exosomes. Interestingly, other exosomal markers such as ALIX, a protein associated with the endosomal sorting complexes required for the transport (ESCRT) machinery during intraluminal vesicle formation, and annexin were not detected. Differences in exosome composition have previously been observed and been suggested to indicate differences in exosome biogenesis^2^.

**Figure 1.**
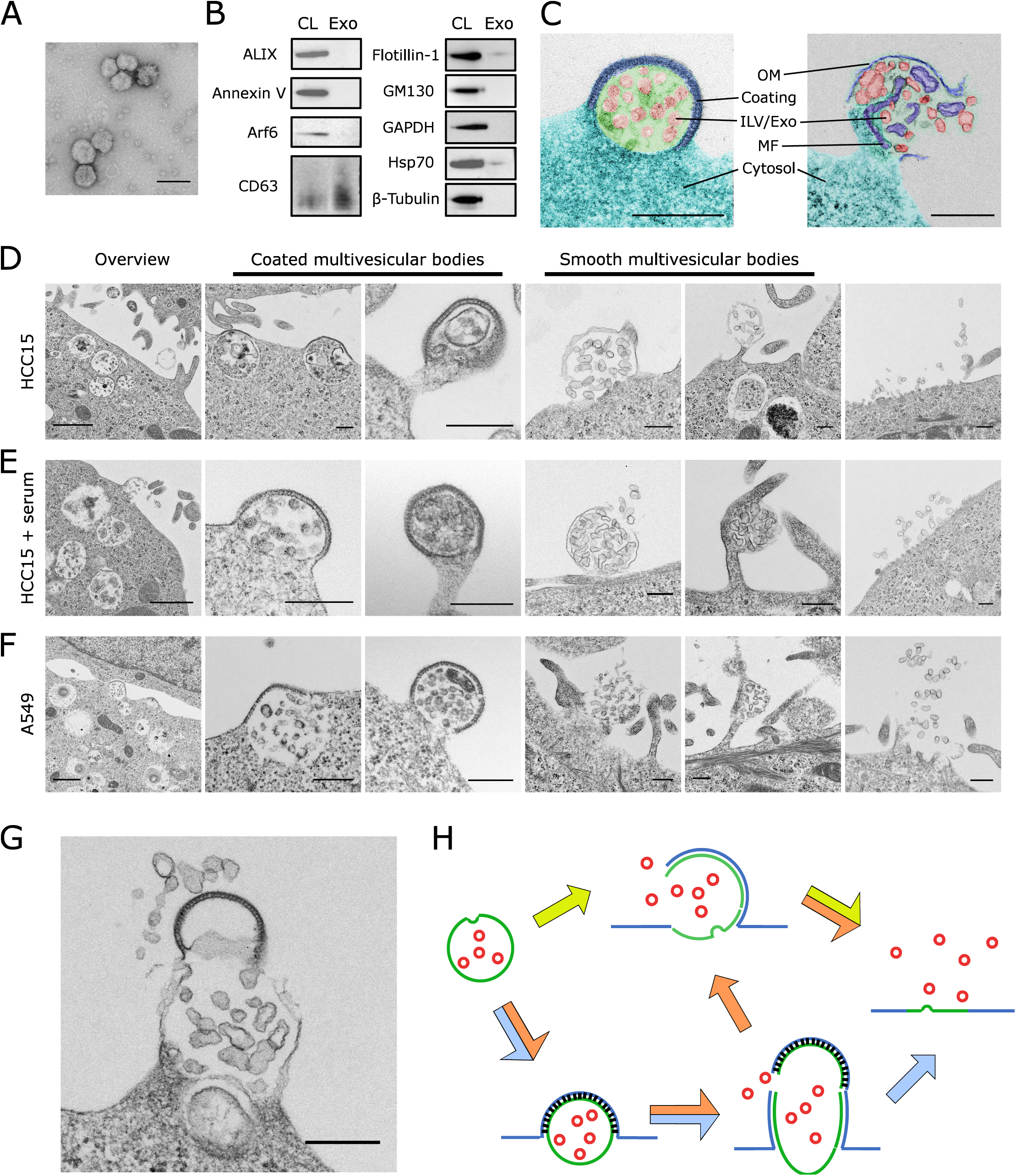
Novel exosome biogenesis routes. (**A**) Morphology of HCC15 exosomes. Negative stain; bar, 100 nm. (**B**) Cellular marker proteins in HCC15 exosomes. CL, cell lysate; Exo, exosome. (**C**) Typical coated (left) and smooth (right) multivesicular structures. Average diameters of the coated and smooth multivesicular structures were 391±71 nm (±*sd, n*=20) and 607±143 nm (*n*=34), respectively. Ultrastructures were highlighted with hue adjustment with the GIMP software. Bar 300 nm; OM, outer membrane; MF, membrane fragments; ILV, intraluminal vesicles. (**D-F**) Cellular ultrastructure related to exosome release. Indicated cells were cultured with (**E**) or without (**D,F**) serum. Bar, 1 µm (overview) or 300 nm (all other images) (**G**) Intermediate structure with characteristics of both coated and smooth multivesicular structures. Cell line, HCC15. Bar, 300 nm. (**H**) Three routes for exosome biogenesis: through the coated (blue arrows) or smooth (green) multivesicular structures, or through both structures consequentially (orange).

Transmission electron microscopy indicated an exosome release pattern that differed from the established membrane fusion model. Instead, the multivesicular structure formed a bulge or blister, sometimes at the end of a long protrusion, which released the exosomes by rupturing (**Fig. 1C-H**). There were two types of multivesicular structures, smooth ones surrounded by an ordinary looking outer membrane and those coated by a thicker electron dense structure with regular spacing patterns (**Fig. 1C).** To our knowledge, such coated multivesicular structures have not been reported before. Both coated and smooth multivesicular structures contained exosome-sized intraluminal vesicles.

The size and morphology of the smooth multivesicular structures (**Fig. 1C**) were similar to that of the multivesicular cargoes/spheresomes/migrasomes released by various cell types^3–5^. However, we did not find multivesicular structures or clustered exosomes detached from the cells. Ruptured smooth multivesicular structures suggested that the intraluminal vesicles were released by a burst and not by fusion with the plasma membrane, which would have resulted in a concave structure as typically described^6^. Both coated and smooth multivesicular structures differed in size (**Fig. 1C**) from known structures that protrude from the membrane, such as larger (>2 µm) entities associated with blebbing and pseudopod formation, and smaller (<0.2 µm) entities associated with virus budding and intracellular vesicle formation in multivesicular endosomes^1,2^. The coated and smooth multivesicular structures may be related to microvesicles, perhaps an immature form^2,7,8^, whose formation mechanism is largely unknown. Further research is required to clarify the exact nature and relationship of these structures.

Exosome release resulted in structurally distinctive areas in the plasma membrane (**Fig. 1D-F**), which were similar to the plasma membrane regions where HIV like particles bud from the cells^9^.

The coated and smooth multivesicular structures as well as the exosome releasing areas of the plasma membrane were also present when the cells were cultured in FBS-containing cell culture medium (**Fig. 1E**) and in A549, another lung cancer cell line^10^ (**Fig. 1F).** In addition, an intermediate structure was found in HCC15 cells (**Fig. 1G**) with an electron dense coating layer, followed with a multivesicular structure, characteristics of both the coated and smooth multivesicular structures. This intermediate structure protruded from the plasma membrane, ruptured and released exosome-sized vesicles.

Based on these discovered structures, we suggest three new routes for exosome biogenesis and release (**Fig. 1H**). 1) Smooth multivesicular structures transport endosomes towards the plasma membrane, protrude and rupture to release exosomes. 2) Coated multivesicular structures carrying endosomes protrude from the plasma membrane and rupture (potentially mechanically) at the base to release exosomes. 3) In intermediate forms, coated membrane areas protrude from smooth multivesicular structures resulting in rupture and exosome release. These three routes seem to coexist in the same cell cultures.

In summary, we discovered exosome biogenesis involving two novel structures, the coated and smooth multivesicular structures, which can bud from the plasma membrane and rupture to release their intraluminal vesicles as exosomes. Altogether, the results point to unique cellular mechanism for membrane association, deformation and fusion. Further investigation is needed to unveil the molecular details that enable and regulate this unconventional process for exosome biogenesis.

## Acknowledgments

This project was supported by the Eastern Academic Research Consortium (Eastern ARC) and MSc programmes Biotechnology and Bioengineering and Cancer Biology and Therapeutics of the School of Biosciences, University of Kent.

