## Supplementary Materials and Methods for "A novel exosome biogenesis mechanism: multivesicular structures budding and rupturing at the plasma membrane"

**Supplementary Information**

**Materials and methods**

**Cell culture**

The non-small cell lung cancer cell lines A549 and HCC15 (DSMZ, Braunschweig, Germany) were cultured at 37°C with 5% CO_2_ in Iscove’s Modified Dulbecco’s Medium (Fisher Scientific) supplemented with 100 I.U./ml penicillin, 100 µg/ml streptomycin, and 10% foetal bovine serum (FBS, Sigma Life Science) unless otherwise specified (see below).

**Exosome preparation with differential centrifugation**

Cells were cultivated in T175 flasks until a confluency of 70-80% was achieved. The cells were washed two times with PBS (Sigma-Aldrich) and continued to be incubated with medium without FBS for 16 h. The medium was then subjected to a series of centrifugation steps: twice at 3 000 g for 10 min (Rotor A-4-81, Centrifuge 5810 R, Eppendorf), once at 10 000 g for 30 min (Rotor Type 70 Ti, Beckman Coulter) and finally once 100 000 g for 60 min (Rotor SW 40 Ti, Beckman Coulter) where the pellets were collected. To reduce their final volume, the 100 000 g pellets, the exosome sample, were re-suspended with PBS, pooled, and re-pelleted again with centrifugation at 196 000 g for 60 min (Rotor SW 40 Ti, Beckman Coulter). The final suspension of the exosome pellets were adjusted to a volume of 150 µl with PBS. The pellets were stored at -80C with PBS before western blot analyses; for negative staining electron microscopy, fresh samples were used. In general, equivalent of 25% total exosomes/pellet collected from one T175 flasks was loaded for western blot analyses.

In parallel, the cellular protein was extracted from the same cells. After the medium was removed for exosome preparation, the cells were washed with cold PBS and lysed with lysis buffer (585 µl, 2% NP-40, 0.2% SDS, 0.5 mM EGTA) supplemented with protease inhibitors (Thermofisher Scientific). After removal of nuclei with centrifugation, the lysate was stored at -80C. A volume of the lysate containing 40 µg protein was used for western blot analyses.

**Western blot analysis**

Western blot analysis was performed in a conventional fashion. In brief, the samples were incubated with non-reducing buffer (25 mM Tris-HCl pH 6.8, 8% SDS, 30% Glycerol, 0.02% Bromophenol Blue) 70°C for 10-15 min before electrophoresis (Novex SDS-PAGE system). After electrophoresis, proteins were transferred (1-Step system, Thermofisher Scientific) onto a nitrocellulose membrane. The membrane was blotted with antibodies in TBST (20 mM Tris, 150 mM NaCl, 0.1% Tween 20) buffer system. The primary or secondary antibodies conjugated with horseradish peroxidase (HRP) were visualised with chemical luminesce (Thermo Scientific/Pierce). The usage of each antibody varied and is listed in the following table.

| **Antibody** | **Source (Catalog #)** | **Dilution** | **Blocking agent: 5% (w/v)** |
| --- | --- | --- | --- |
| CD63-HRP | Santa Cruz Biotechnology (MC-49.129.5) | 1:1000 | Non-fat dry milk |
| Arf6 | Cell Signalling Technology (D12G6) | 1:1000 | BSA |
| GAPDH-HRP | Cell Signalling Technology (14C10) | 1:1000 | Non-fat dry milk |
| ALIX | Cell Signalling Technology (2171) | 1:1000 | Non-fat dry milk |
| Annexin V | Cell Signalling Technology (8555) | 1:1000 | BSA |
| Hsp70 | Cell Signalling Technology (4876) | 1:1000 | BSA |
| Flotillin-1 | Cell Signalling Technology (18634) | 1:1000 | BSA |
| β-Tubulin-HRP | Cell Signalling Technology (2146S) | 1:1000 | Non-fat dry milk |
| Anti-mouse IgG-HRP | Cell Signalling Technology (7076) | 1:500 | Non-fat dry milk |
| Anti-rabbit IgG-HRP | Cell Signalling Technology ( 7074S) | 1:500 | Non-fat dry milk |

**Transmission electron microscopy (TEM)**

Exosomes (5 µL) were applied to 600 mesh formvar/carbon coated grids and allowed to settle down for 5 minutes. The sample was then fixed in situ by adding 5 µl of 5% glutaraldehyde in 200mM sodium cacodylate (CAB) buffer pH7.2 for 5 minutes before being transferred into a 10 µl drop of 2.5% glutaraldehyde in CAB for 10 minutes. Grids were then washed by passing through 3 x 10 µl drops of Milli Q water 2 minutes in each drop. Grids were then dried and negatively stained with 2% aqueous Uranyl acetate and were viewed in a Jeol 1230 TEM at an accelerating voltage of 80 kV equipped with a Gatan One View camera.

Cells were grown on UV-sterilized Aclar film (Agar Scientific) in 24 well plates (Greiner) until a confluency of 70-80% was achieved. When indicated (with serum), the cells were kept in the same medium (with FBS); under any other circumstances, the cells were washed with PBS and the medium was exchanged to FBS-free fresh medium. After the cells were further cultivated for 16 h, the medium was removed and the cells were fixed *in situ* in wells of 24 well plates for 0.5h with 2.5% glutaraldehyde in 100 mM CAB buffer pH 7.2. The Aclar with cells was then transferred to a 7 ml glass vial in the same fixative for a further hour. Samples were then washed in 100 mM CAB 2 x 10 minutes and were then post fixed with 1% Osmium tetroxide in 100 mM CAB for 1 h. Samples were washed in Milli Q water 2x10 minutes. Samples were then placed in 50% ethanol for 10 minutes and then in 70% ethanol over night at 4°C. The next morning dehydration was completed 90% ethanol 10 minutes and 3 x 10 minutes 100% dry ethanol. Samples were then placed in Propylene oxide 2x 10 minutes. Samples were then placed in 1:1 Propylene oxide: Agar LV resin (Agar Scientific) for 30 minutes. Samples were then placed in pure Agar LV resin 2 x 1.5 h and were then placed in 6 ml aluminium dishes (cell side up) and were polymerised at 60°C for 24 h. When polymerised, approximately one quarter of the cells on Aclar was cut out with a jig saw and superglued onto a blank resin block. The Aclar was then peeled off leaving the cells in the surface of the resin block. Sections of 70 nm were then cut through the plane of the cells using a Leica EM UC7 ultramicrotome equipped with a Diatome diamond knife. Sections were picked up on 400 mesh uncoated copper grids (Agar Scientific) and allowed to air dry. Sections on grids were then counterstained in 4.5% Uranyl acetate in 1% acetic acid for 45 minutes, then washed for 10 seconds in a stream of Milli Q water from a wash bottle. This was followed by 7 minutes in Reynolds lead citrate and then washed in Milli Q water and dried. Sections were viewed in a Jeol 1230 TEM equipped with a Gatan One View camera at an accelerating voltage of 80 kV.
